# Neuronal alterations in AKT isotype expression in schizophrenia

**DOI:** 10.1101/2023.06.19.545479

**Authors:** Emily A. Devine, Alex W. Joyce, Ali S. Imami, Abdul-rizaq Hammoud, Hasti Golchin, Hunter Eby, Elizabeth A. Shedroff, Sophie M. Asah, Consuelo Walss-Bass, Sinead O’Donovan, Robert E. McCullumsmith

## Abstract

Schizophrenia is characterized by substantial alterations in brain function, and previous studies suggest insulin signaling pathways, particularly involving AKT, are implicated in the pathophysiology of the disorder. This study demonstrates elevated mRNA expression of AKT1-3 in neurons from schizophrenia subjects, contrary to unchanged or diminished total AKT protein expression reported in previous postmortem studies, suggesting a potential decoupling of transcript and protein levels. Sex-specific differential AKT activity was observed, indicating divergent roles in males and females with schizophrenia. Alongside AKT, upregulation of PDK1, a critical component of the insulin signaling pathway, and several protein phosphatases known to regulate AKT were detected. Moreover, enhanced expression of the transcription factor FOXO1, a regulator of glucose metabolism, hints at possible compensatory mechanisms related to insulin signaling dysregulation. Findings were largely independent of antipsychotic medication use, suggesting inherent alterations in schizophrenia. These results highlight the significance of AKT and related signaling pathways in schizophrenia, proposing that these changes might represent a compensatory response to a primary defect of insulin signaling. This research underscores the need for a detailed understanding of these signaling pathways for the development of effective therapeutic strategies.

## Introduction

AKT, named after the AK mouse strain plus “transforming” or “thymoma” (also known as Protein Kinase B), is a “hub” serine/threonine protein kinase involved in the regulation of cellular processes including cell growth, proliferation, differentiation, migration, survival, and metabolism [1]. The highly conserved AKT kinase family consists of three isotypes, AKT1, AKT2, and AKT3, each encoded by separate genes [2]. Although previously thought to be functionally redundant, recent work suggests these highly homologous AKT isotypes exhibit diverse and unique functions [3]. Isotype-specific functions were demonstrated using single and double AKT isotype-specific knockout mice. Knockout studies suggest associations with AKT1 for whole organismal growth and survival, AKT2 with glucose metabolism, and AKT3 with brain development and size [3].

Dysregulation of the AKT signaling pathway is associated with schizophrenia, a severe neuropsychiatric disorder [4–6]. One of the key characteristics of schizophrenia is cognitive deficits, notably in attention, executive function, and memory [7, 8], all of which are regulated by AKT and its canonical signaling pathways [9, 10]. Disruptions in glucose metabolism, another mechanism regulated by AKT [11], are also associated with cognitive deficits [12, 13]. Further, isotype-specific knockouts have schizophreniform endophenotypes, with AKT1 and AKT3-deficient mice exhibiting deficits in social behaviors, learning, and memory [14, 15]. In humans, initial genome-wide association studies (GWAS) found schizophrenia risk with genetic variations in AKT1, and more recently AKT3 [5, 16, 17].

Alterations in AKT and associated signaling pathways have also been explored in postmortem brain in persons with schizophrenia, with several studies showing reduced AKT1 protein expression [5, 16, 18–21]. Further, we previously identified a protein kinase network associated with the pathophysiology of schizophrenia where alterations in AKT signaling were suggested as an aberrant signaling node [22, 23]. These alterations were confirmed with pan-AKT western blots and enzyme activity assays showing a decrease in phospho-AKT protein expression and an increase in AKT-specific activity in the anterior cingulate cortex (ACC)[23]. Taken together, these findings provide robust evidence for alterations of AKT signaling pathways in the pathophysiology of schizophrenia [5].

While these previous studies are informative, investigating specific AKT isotypes at the cellular level is an important next step. Region-level brain studies suffer from the “blender problem,” where all cell types are mixed in homogenized brain samples. Studies using blended brain samples may be difficult to interpret, as measures of gene expression or enzyme activity will reflect the aggregate of changes across cell types; if expression goes up in one cell type, and down in another, the net effect may be no change [24, 25]. Further, AKT has cell-subtype-specific functions, particularly during development [4, 9, 26]. Finally, prior work has also implicated schizophrenia-associated genetic variations of AKT with disruptions in cortical neuron signaling, leading to the cognitive deficits present in the disease [27]. Taken together, these observations support cell-level investigation of AKT isoform expression in neurons.

In the present study, we examine transcript expression for genes in the AKT signaling pathways in anterior cortex (ACC) pyramidal neurons in schizophrenia. We focused on pyramidal neurons due to the convergence of genetic risk in schizophrenia on glutamatergic synapses; We focused on the ACC due to its roles in cognition and executive function, which are often impaired in this illness [28, 29]. First, we coupled laser microdissection (LMD) with QPCR to establish changes in AKT isotypes and associated pathway components in pyramidal neurons at the transcript-level. Second, we investigated the potential effects of schizophrenia-linked single nucleotide polymorphisms (SNPs) of AKT on gene expression. Lastly, we reanalyzed our published ACC kinase activity array dataset [23] using recombinant AKT activity profiles to assess changes in AKT1 and AKT3 activity in schizophrenia.

## Materials and Methods

### Subjects

Postmortem tissue used in regional level and cell-level gene expression and genotyping studies will be referred to as “cohort 1,” while the postmortem cohort we reassessed from a previously published study will be referred to as “cohort 2.” Anterior cingulate cortex (ACC) tissue for both cohorts was obtained from the Mount Sinai NIH Brain and Tissue Repository (New York, New York). Schizophrenia (n = 20 for cohort 1, n = 12 for cohort 2) and control (n = 20 for cohort 1, n = 12 for cohort 2) subjects were matched for age, sex, postmortem interval (PMI), and tissue pH (Tables 1 and 2). The two cohorts do not overlap. Brains were freshly frozen and stored at −80°C until needed for studies. For the region-level gene expression and genotyping studies, tissue was sectioned into 14 μm thick sections on SuperfrostPlus glass slides (Fisher Scientific, Waltham, MA). The tissue used for cell-level gene expression studies was cryostat sectioned into 12 μm thick sections on PEN membrane glass slides (Leica Microsystems, Wetzlar, Germany).

**Table 1.** Cohort 1 Subject Demographics

| COHORT 1 DX | N | SEX | PMI | AGE | PH | RIN | ANTIPSYCHOTICS<br>(ON/OFF/UNKNOWN) |
| --- | --- | --- | --- | --- | --- | --- | --- |
| CONTROL | 20 | 7F/13M | 12.4±7.5 | 78±7 | 6.6±0.4 | 4.4±1.9 | N/A |
| SCHIZOPHRENIA | 20 | 9F/11M | 13.1±5.8 | 75±8 | 6.2±0.3 | 4.1±1.6 | 12/6/2 |

Demographics of Mount Sinai NIH Brain and Tissue Repository subjects used in Cohort 1. Dx, diagnosis; N, number of subjects; F, female; M, male; PMI, postmortem interval (hours); N/A, not applicable. Data presented as mean +/- standard deviation.

**Table 2.** Cohort 2 Subject Demographics

| COHORT 2 DX | N | SEX | PMI | AGE | PH | ANTIPSYCHOTICS<br>(ON/OFF/UNKNOWN) |
| --- | --- | --- | --- | --- | --- | --- |
| CONTROL | 12 | 6 F/6 M | 9.1±6.2 | 75±10.6 | 6.4±0.2 | N/A |
| SCHIZOPHRENIA | 12 | 6 F/6 M | 10.7±4.6 | 75±11.7 | 6.2±0.3 | 9/2/1 |

Demographics of Mount Sinai NIH Brain and Tissue Repository subjects used in Cohort 2. DX, diagnosis; N, number of subjects; F, female; M, male; PMI, postmortem interval (hours); RIN, RNA integrity number; N/A, not applicable. Data presented as mean +/- standard deviation.

### Laser Microdissection

The LMD6 (Leica Microsystems) was used for laser microdissection (LMD). Frozen tissue sections from cohort 1 were thawed at room temperature and allowed to dry. Tissue sections were then rehydrated with RNase-Free H^2^O and were nissl stained with an RNAse-free cresyl-violet solution (FD NeuroTechnologies, Columbia, MD). Slides were then dehydrated through serial ethanol washes. Enriched populations of pyramidal neurons (500 per subject) were identified via morphology and cut from the gray matter of the ACC at an objective lens of 40x as described in our previously validated protocol [30–34]. The laser settings consisted of power: 24-25, aperture: 4-5, and speed: 8. The dissected cells were collected into the cap of separate 0.5 mL tubes (Axygen, Union City, CA) for each subject and incubated with 30 μL of PicoPure RNA extraction buffer (Applied Biosystems, Foster City, CA) for 32 min at 40°C. Samples were then centrifuged for 2 min at 400 × g and stored at −80°C.

### RNA Isolation and Reverse Transcription

For the region-level study, RNA was extracted from cryosections of the ACC using the RNeasy Minikit (Qiagen, Hilden, Germany) following the manufacturer’s protocol. RNA concentration was measured via nanodrop and all subjects were normalized to 6.4 ng/μL. For the cell-level study, RNA was isolated from laser microdissected pyramidal neurons using the PicoPure RNA isolation kit (Applied Biosystems) according to the manufacturer’s protocol. In both the region and cell-level studies, cDNA was synthesized using the High-Capacity cDNA Reverse Transcription Kit (Applied Biosystems). After reverse transcription region-level cDNA was diluted 1:3 with RNase/DNase-free water and stored at −20°C until used in RT-PCR. Pyramidal neuron cDNA was then pre-amplified.

### cDNA Pre-Amplification

Taqman assays (Supplementary Table 1) were pooled and diluted with RNase/DNase Free water to a final concentration of 0.2x and were combined with Fast Start Universal Master Mix (Roche, Basel, Switzerland) and cDNA for the preamplification PCR reaction. The PCR cycles were: 1 cycle held at 95°C for 10 min, then 14 cycles of denaturing at 95°C for 15 sec and annealing at 60°C for 1 min. Pre-amplified samples were diluted 1:5 with RNase/DNase-free water and stored at 20°C until used in RT-PCR.

### Quantitative RT-PCR

In the region and cell-level studies, RT-PCR TaqMan gene expression assays (Applied Biosystems) were used to measure the expression of β-actin (ACTB), cyclophilin A (PPIA), glyceraldehyde-3-phosphate dehydrogenase (GAPDH), beta-2-microglobulin (B2M), AKT serine/threonine kinase 1 (AKT1), AKT serine/threonine kinase 2 (AKT2), AKT serine/threonine kinase 3 (AKT3), forkhead box O1 (FOXO1), Phosphatase and tensin homolog (PTEN), PH domain and leucine-rich repeat protein phosphatase 1 (PHLPP1), PH domain and leucine-rich repeat protein phosphatase 2 (PHLPP2), 3-phosphoinositide dependent protein kinase 1 (PDPK1), and protein phosphatase 2 catalytic subunit alpha (PPP2CA). Each reaction was performed in duplicate in a 20 μL volume consisting of 10 μL Fast Start Universal Master Mix (Roche), 1 μL TaqMan probe, 6 μL RNAse/DNAse free water, and 3 μL cDNA. Cycling conditions included a 10-minute hold at 95°C followed by 40 cycles of 95°C for 15 seconds and 60°C for 1 minute. A pooled calibrator cDNA sample and a set of serial dilutions ranging from 1:5 to 1:40 in regionlevel studies and 1:5 to 1:80 in cell-level studies were included on each plate to determine a standard curve for the quantification of each gene. After making up the pooled cDNA sample, samples were further diluted 1:2, with the exception of subjects 24508 and 23847 which were further diluted 1:4 in the cell-level studies. This was because these subjects had significantly higher RNA concentrations than the others and we cannot normalize RNA concentrations at the cellular level due to their small amounts of RNA present. For the negative controls, cDNA was replaced with an equivalent volume of RNAse/DNAse-free water. Each assay was performed in 96-well optical reaction plates (Applied Biosystems) on an ABI Stepone Plus (Applied Biosystems) qPCR system. The standard curve method was used for relative quantification. The values for duplicate samples were averaged and normalized to the geometric mean of housekeeping genes PPIA, GAPDH, and B2M for the region-level study and additionally with ACTB in the cell-level study.

### Single Nucleotide Polymorphism Study

gDNA was extracted from cryosections of the ACC with the QIAMP DNA Mini Kit (Qiagen) according to the manufacturer’s protocol. DNA concentration was measured via nanodrop and all subjects were normalized to 16.2 ng/uL. Taqman genotyping assays (Applied Biosystems) for AKT SNPs rs1130214 (C_26352825_10), rs2494732 (C_16191608_10), rs1130233 (C_7489835_10), and rs3730358 (C_193157_10) were used to genotype ACC samples. Each reaction was performed in duplicate in a 25 μL volume consisting of 12.5 μL Fast Start Universal Master Mix (Roche), 1.25 μL TaqMan assay working stock (20x), and 11.25 μL gDNA. Cycling conditions included a 10-minute hold at 95°C followed by 40 cycles of 95°C for 15 seconds and 60°C for 1 minute. Each assay was performed in 96-well optical reaction plates (Applied Biosystems) on an ABI Stepone Plus (Applied Biosystems) qPCR system.

### Antipsychotic Analysis

To determine whether chronic treatment of antipsychotics effect our findings, we performed *in silico* analyses (also called “lookup studies”) using Kaleidoscope (https://cdrl.shinyapps.io/Kaleidoscope/), an R shiny application that contains publicly available omics datasets from psychiatric disorders, as well as pharmacological treatment studies in models systems [35]. Our genes of interest (AKT1, AKT2, AKT3, FOXO1, PTEN, PPHLP1, PPHLP2, PDPK1, PPP2CA) were “looked up” in pharmacological databases with diverse substrates and treatments, including typical and atypical antipsychotics medications. Databases were separated into typical and atypical antipsychotic groups and these two groups were further separated into the groups, “frontal cortex” and “other brain regions.” Tables 2-5 list the number and percent of significant results (P ≤ 0.05) as well as the average Log Fold Change for each gene within these groups. Fold Change, Log Fold Change, and P-values for our genes of interest from each database can be found in an excel file in the supplementary materials.

### Kinome Array Profiling

Recombinant AKT1, AKT2, and AKT3 were purchased from ReactionBiology and run individually on the Pamgene 12 kinome array serine/threonine kinase (STK) chip. Activity profiles for reporter peptides were analyzed as previously described using Bionavigator software [22, 23, 36–39]. Peptides with fold-change > +/-15% were carried forward for subsequent analyses. Principle component analysis (PCA) was used to cluster peptides associated with higher, middle, and low levels of recombinant AKT1 and AKT3 protein kinase activity on the array. Recombinant AKT2 did not give a measurable signal on the array and was not further considered. Pathway analyses for “high-affinity” peptides were performed using EnrichR with the BioPlanet2019 database. Fold change values for high-affinity AKT1 and AKT3 peptides for subject pairs from our previously published postmortem ACC schizophrenia study were extracted from the original datasets and displayed as log2 fold change (for each peptide) for 12 of the subject pairs in this study. We also reassessed the activity for the AKT1-and AKT3-reporting peptides in a chronic haloperidol treatment dataset from the kinome array. Rats were treated for 9 months with 28.5 mg/kg haloperidol-decanoate or vehicle (sesame oil), and the frontal pole was assessed on the STK chip [23]. Fold change values for high-affinity AKT1 and AKT3-specific peptides for our previously published haloperidol kinome array study were extracted from the original datasets and displayed as log2 fold change (for each peptide) for haloperidol and control groups in this study.

### Data Analysis

Alpha = 0.05 for all statistical tests. Data were analyzed with Statistica (TIBCO Software, Palo Alto, CA) and GraphPad Prism 9 (GraphPad Software, La Jolla, CA). All data sets were tested for normal distribution (D’Agostino and Pearson omnibus normality test) and homogeneity of variance (F-test). Outliers were excluded using the ROUT method with Q set to 1%.

#### Quantitative RT-PCR and SNP Assays

Data were log-transformed. Correlation analyses were performed to determine associations between transcript expression and age, PMI, and RIN value. Analysis of covariance (ANCOVA) was performed if significant correlations were found. If no correlations were present, data were analyzed with Student’s t-test, Welch’s t-test, or Mann-Whitney test.

## Results

The mRNA expression levels of the AKT serine/threonine kinase isotypes (AKT1, AKT2, AKT3) and components of their signaling pathway (FOXO1, PTEN, PHLPP1, PHLPP2, PDPK1, PPP2CA) were measured in tissue homogenates (region-level studies) and enriched populations of pyramidal neurons (cell-level studies) in the ACC in schizophrenia in cohort 1.

### Region-Level Gene Expression Studies

In the tissue homogenates of the ACC, there was a significant increase in mRNA expression in schizophrenia subjects compared to controls for AKT1 (p=0.005), AKT2 (p=0.006), AKT3 (p=0.043), FOXO1 (p=0.033), PTEN (p=0.007), PHLPP2 (p= <0.0001), PDPK1 (p=0.001), and PPP2CA (p=0.02). There was no change detected in mRNA expression for PHLPP1 (p=0.188). There were no significant associations for mRNA expression between AKT1, AKT2, AKT3, FOXO1, PTEN, PHLPP1, PHLPP2, PDPK1, PPP2CA, and age, pH, PMI or RIN values (Figure 1).

**Figure 1.**
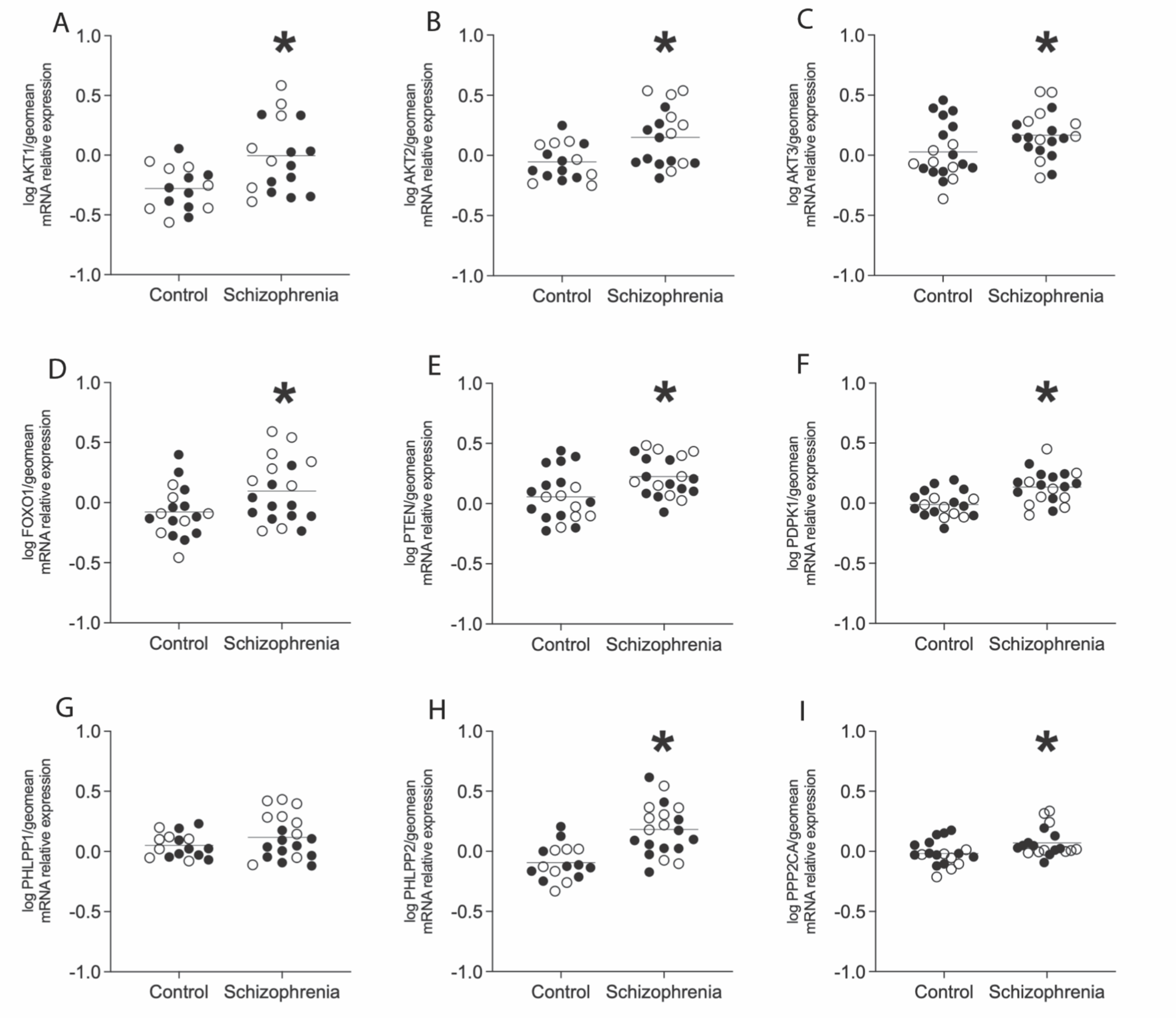
Region-level expression of AKT serine/threonine protein kinase isotypes and pathway components. Open circles indicate females, closed circles indicate males. Analysis revealed increased expression of A) AKT1, B) AKT2, C) AKT3, D) FOXO1, E) PTEN, F) PDPK1, H) PHLPP2, and I) PPP2CA (*p<0.05). There was no significant change in gene expression of G) PHLPP1 in schizophrenia subjects compared to controls. Data are log-transformed and analyzed using either Student’s t-test or Welch’s t-test. Data expressed as mean +/-SEM, n=15-20/group. Abbreviations: AKT serine/threonine kinase 1 (AKT1), AKT serine/threonine kinase 2 (AKT2), AKT serine/threonine kinase 3 (AKT3), Forkhead box O1 (FOXO1), Phosphatase and tensin homolog (PTEN), 3-phosphoinositide dependent protein kinase 1 (PDPK1), PH domain and leucine-rich repeat protein phosphatase 1 (PHLPP1), PH domain and leucine-rich repeat protein phosphatase 2 (PHLPP2), Protein phosphatase 2 catalytic subunit alpha (PPP2CA).

### Cell-Level Gene Expression Studies

In a population of enriched pyramidal neurons from the ACC, there was a significant increase in mRNA expression in schizophrenia subjects compared to controls for AKT1 (p=0.002), AKT2 (p=0.008), AKT3 (p=0.006), FOXO1 (p=0.021), PTEN (p=0.009), PHLPP2 (p=0.019), and PDPK1 (p=0.01). There were no changes detected in mRNA expression for PHLPP1 (p=0.928), or PPP2CA (p= >0.999). There were no significant associations for mRNA expression between AKT1, AKT2, AKT3, FOXO1, PTEN, PHLPP1, PHLPP2, PDPK1, PPP2CA, and age, pH, PMI or RIN values (Figure 2).

**Figure 2.**
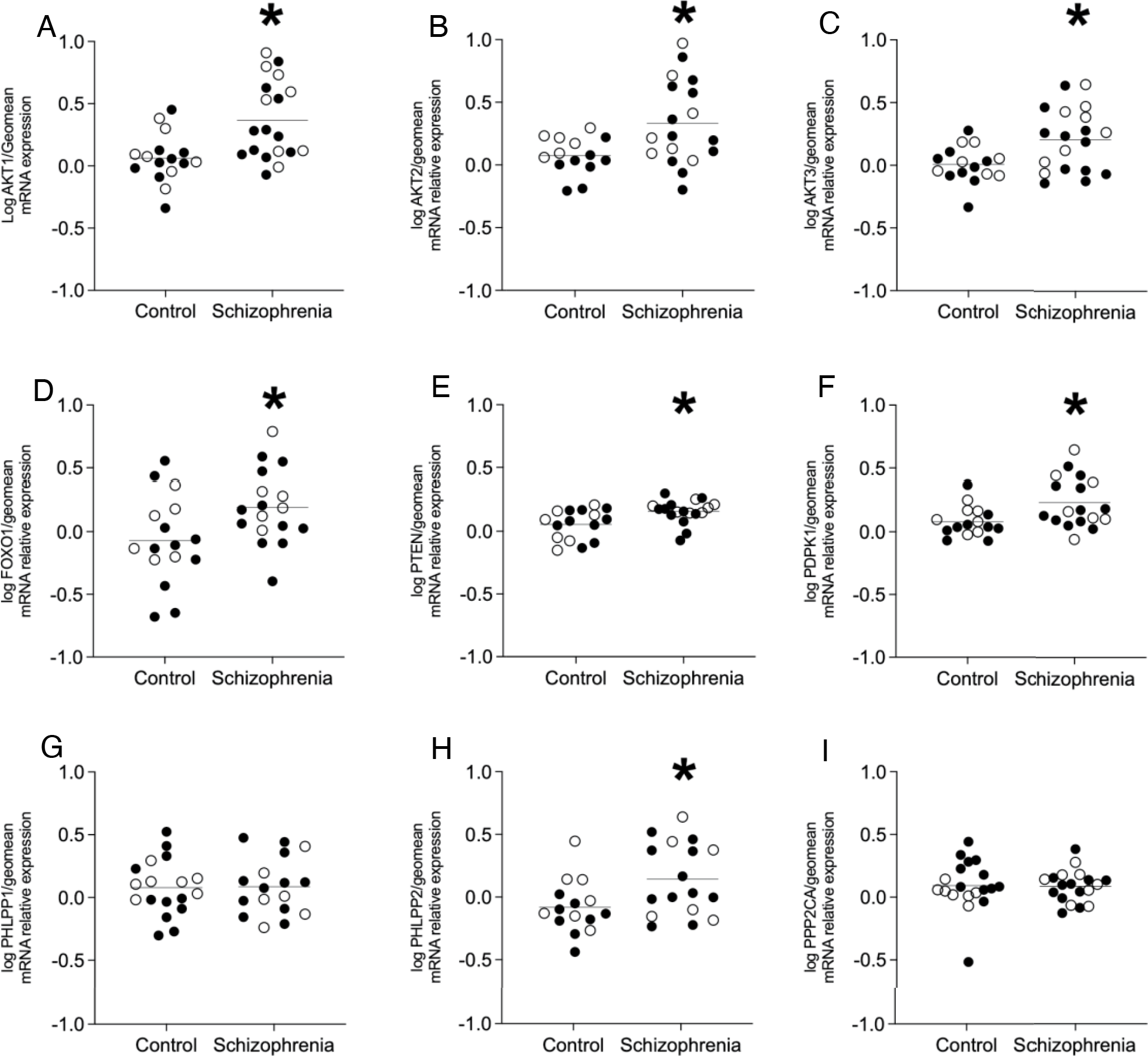
Enriched pyramidal cell population expression of AKT serine/threonine protein kinase isotypes and pathway components. Open circles indicate females, closed circles indicate males. Analysis revealed increased expression of A) AKT1, B) AKT2, C) AKT3, D) FOXO1, E) PTEN, F) PDPK1 and H) PHLPP2 (*p<0.05). There was no significant change in gene expression of G) PHLPP1 or I) PPP2CA in schizophrenia subjects compared to controls. Data are log-transformed and analyzed using either Student’s t-test, Welch’s t-test, or Mann-Whitney test. Data mean+/-SEM, n=14-19/group. AKT1 AKT serine/threonine kinase 1, AKT2, AKT serine/threonine kinase 2, AKT3 AKT serine/threonine kinase 3, FOXO1 Forkhead box O1, PTEN Phosphatase and tensin homolog, PDPK1 3-phosphoinositide dependent protein kinase 1, PHLPP1 PH domain and leucine-rich repeat protein phosphatase 1, PHLPP2 PH domain and leucine-rich repeat protein phosphatase 2, PPP2CA, Protein phosphatase 2 catalytic subunit alpha.

### Effects of Sex

We found increased AKT1 and AKT3, but not AKT2, in pyramidal neurons in female subjects with schizophrenia (Fig S1). No changes were detected in male subjects.

### Genotyping Single Nucleotide Polymorphism Studies

The ACC tissue homogenate of the schizophrenia (n=20) and control subjects was genotyped for AKT1 SNPs rs1130214, rs2494732, rs1130233, and rs3730358. These SNPs were chosen due to their association with the inheritance of schizophrenia and their role in impaired cognition [27]. AKT1 gene expression was analyzed by genotype regardless of the subject’s diagnosis. For SNPs rs3730358 (p=0.08), rs2494732 (p=0.06), rs1130233 (p=0.9), and rs1130214 (p=0.45) there were no significant differences in gene expression detected (Figure 3).

**Figure 3.**
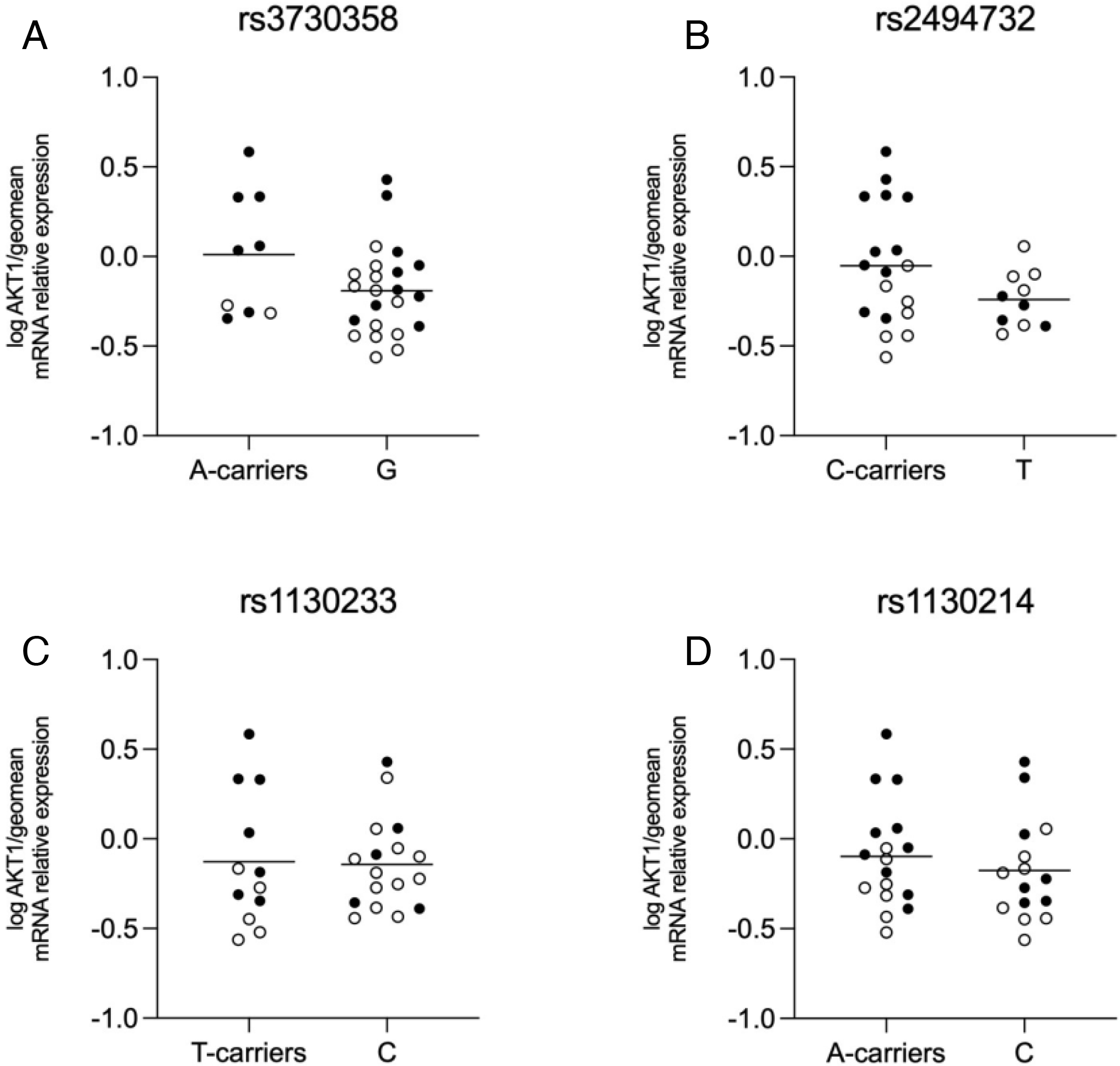
mRNA expression of AKT1 SNPs. Open circles indicate control subjects, and closed circles indicate schizophrenia subjects. There was no significant difference in expression in A) SNP rs3730358 when comparing A-carriers to the G/G polymorphism, B) SNP rs2494732 when comparing C-carriers to the T/T polymorphism, C) SNP rs1130233 when comparing T-carriers to the C/C polymorphism, and D) SNP rs1130214 when comparing A-carriers to the C/C polymorphism. Data are log-transformed and analyzed using either Student’s t-test or Welch’s t-test. Data mean+/-SEM, n=9-23/group. AKT1 AKT serine/threonine kinase 1, SNP single nucleotide polymorphism.

### Effects of Antipsychotic Drugs

To examine the potential effect of chronic antipsychotic treatment, *in silico* analyses were run on over 50 reposited pharmacological datasets that explored transcript expression changes for our genes of interest in antipsychotic-treated rats, mice, and/or hiPSCs. The datasets were divided and analyzed as 4 separate groups and results are summarized in their respective tables: typical antipsychotics in the frontal cortex (Table 3) and other brain regions (Table 4), as well as atypical antipsychotics in the frontal cortex (Table 5) and other brain regions (Table 6). Typical antipsychotics in the frontal cortex showed no changes in AKT1, PHLPP2, and PPP2CA gene expression in any of the datasets, while AKT2, AKT3, PDPK1, and PTEN showed decreased mRNA expression, while FOXO1 and PHLPP1 were increased. Atypical antipsychotics in the frontal cortex showed no changes in AKT1, AKT2, PHLPP1, PHLPP2, and PPP2CA in any of the datasets. For AKT3 and PTEN, atypical antipsychotics in the frontal cortex datasets showed divergent results, while PDPK1 gene expression was decreased. Finally, one dataset showed decreased FOXO1 mRNA expression for atypical antipsychotics in the frontal cortex. Data for individual datasets are provided in the supplementary materials.

**Table 3.** Effects of Typical Antipsychotic Medications in the Frontal Cortex

|  | AKT1 | AKT2 | AKT3 | FOXO1 | PTEN | PHLPP1 | PHLPP2 | PDPK1 | PPP2CA |
| --- | --- | --- | --- | --- | --- | --- | --- | --- | --- |
| TOTAL DATASETS | 6 | 6 | 6 | 7 | 7 | 6 | 2 | 7 | 7 |
| SIGNIFICANT DATASETS | 0<br>(0%) | 1<br>(16.7%) | 1<br>(16.7%) | 2<br>(28.6%) | 1<br>(14.3%) | 1<br>(16.7%) | 0<br>(0%) | 3<br>(42.9%) | 0<br>(0%) |
| AVERAGE LOG FOLD CHANGE | N/A | -0.18 | -0.22 | 0.6 | -0.24 | 0.22 | N/A | -0.25 | N/A |

Summary of in silico studies from online databases comparing frontal cortex gene expression in rats, mice, and hiPSCs treated with typical antipsychotics and controls. Significant datasets

**Table 4.**
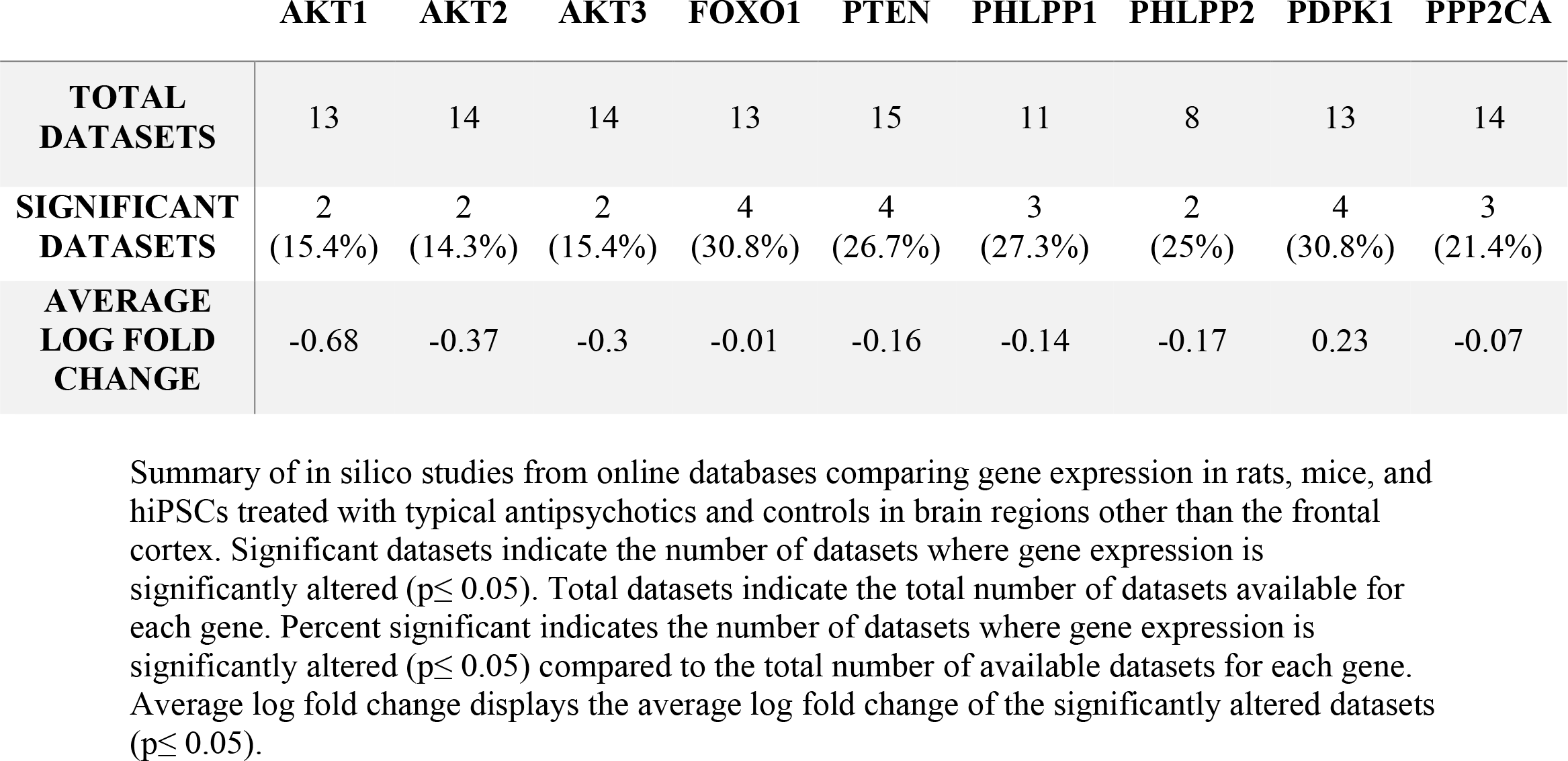
Effects of Typical Antipsychotic Medications in Other Brain Regions

**Table 5.** Effects of Atypical Antipsychotics in the Frontal Cortex

|  | AKT1 | AKT2 | AKT3 | FOXO1 | PTEN | PHLPP1 | PHLPP2 | PDPK1 | PPP2CA |
| --- | --- | --- | --- | --- | --- | --- | --- | --- | --- |
| **TOTAL DATASETS** | 9 | 9 | 9 | 9 | 9 | 8 | 5 | 8 | 9 |
| **SIGNIFICANT DATASETS** | 0<br>(0%) | 0<br>(0%) | 2<br>(22.2%) | 1<br>(11.1%) | 2<br>(22.2%) | 0<br>(0%) | 0<br>(0%) | 1<br>(12.5%) | 0<br>(0%) |
| **AVERAGE LOG FOLD CHANGE** | N/A | N/A | 0.04 | -1.06 | -0.03 | N/A | N/A | -0.18 | N/A |

Summary of in silico studies from online databases comparing frontal cortex gene expression in rats, mice, and hiPSCs treated with atypical antipsychotics and controls. Significant datasets indicate the number of datasets where gene expression is significantly altered ( $p \leq 0.05$ ). Total datasets indicate the total number of datasets available for each gene. Percent significant

**Table 6.** Effects of Atypical Antipsychotics in Other Brain Regions.

|  | **AKT1** | **AKT2** | **AKT3** | **FOXO1** | **PTEN** | **PHLPP1** | **PHLPP2** | **PDPK1** | **PPP2CA** |
| --- | --- | --- | --- | --- | --- | --- | --- | --- | --- |
| **TOTAL DATASETS** | 34 | 34 | 35 | 36 | 36 | 29 | 27 | 36 | 33 |
| **SIGNIFICANT DATASETS** | 9<br>(26.5%) | 10<br>(29.4%) | 4<br>(11.4%) | 16<br>(44.4%) | 10<br>(27.8%) | 8<br>(27.6%) | 6<br>(22.2%) | 12<br>(33.3%) | 3<br>(9.1%) |
| **AVERAGE LOG FOLD CHANGE** | 0.01 | -0.06 | -0.02 | -0.07 | -0.36 | -0.02 | 0.01 | -0.10 | -0.03 |

Summary of in silico studies from online databases comparing gene expression in rats, mice, and hiPSCs treated with atypical antipsychotics and controls in brain regions other than the frontal cortex. Significant datasets indicate the number of datasets where gene expression is significantly altered ( $p \leq 0.05$ ). Total datasets indicate the total number of datasets available for each gene. Percent significant indicates the number of datasets where gene expression is significantly altered ( $p \leq 0.05$ ) compared to the total number of available datasets for each gene. Average log fold change displays the average log fold change of the significantly altered datasets ( $p \leq 0.05$ ).

### Recombinant Kinase Analysis

We have previously reported decreased phospho-AKT expression and increased specific activity in ACC tissue homogenate in schizophrenia [23]. In these same samples we also previously performed kinase activity array analyses showing AKT as a possible perturbed node in schizophrenia. Since the assignment of upstream kinases for this original analysis was performed via mapping by in silico databases, we assessed recombinant AKT1 and AKT3 protein activity on the same kinase activity array to empirically determine the reporter peptides that best report AKT1 or AKT3 activity (Fig 4A, B). High, medium, and low-affinity peptides were identified using principal component analyses (Fig 4C, D). The top pathways associated with the proteins containing the high-affinity peptides from the array were determined using EnrichR with the BioPlanet2019 database (Fig 4E, F). Consensus peptide sequences were generated for high, medium, and low-affinity peptides (Fig S2). Log2 fold-change expression of kinase activity in region-level ACC brain homogenate was determined for female (left panel) and male subject pairs (Fig 5A, C) for each high-affinity reporter peptide (red and black circles). We then reanalyzed another published dataset [23] using the same approach to determine changes in the high, medium, and low-affinity peptides (red and black circles) in haloperidol-treated rats (28.5mg/kg haloperidol-decanoate every three weeks for 9 months) versus vehicle (sesame oil)(Fig 5B, D). For the human experiment (Fig 5A, C), 12 pairs of subjects were run case-control (ie not pooled), while for the rodent experiment (Fig B, D), n = 10 animals per group were pooled for the kinase activity array.

**Figure 4.**
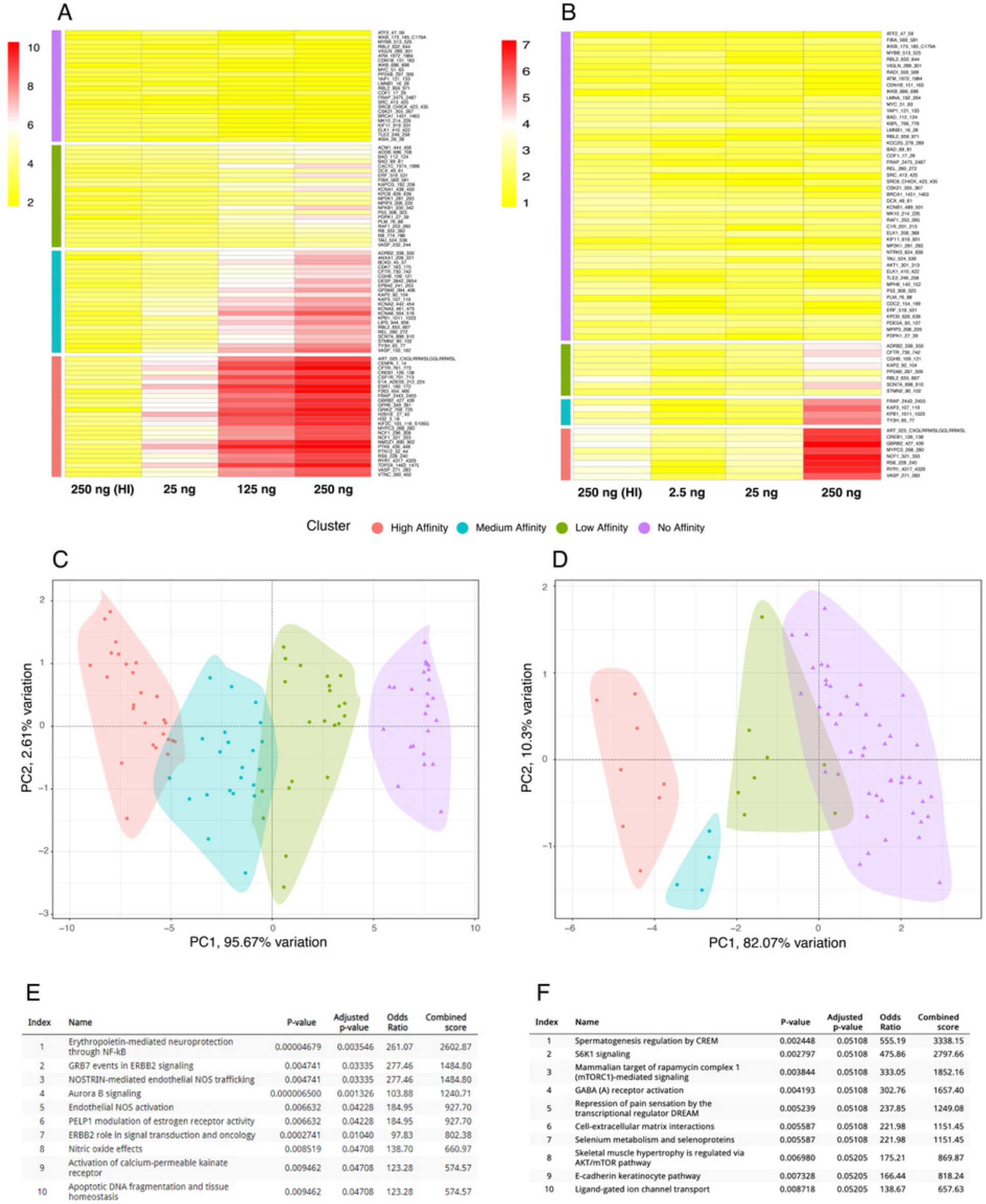
Recombinant AKT kinome array profiling. Recombinant AKT1 or AKT3 (25-250ng) was run in triplicate on the PamGene kinome array, along with 250ug of heat-inactivated (95C x 10 minutes) recombinant AKT1 or AKT3 protein as a negative control (panels A and B). Principle component analysis of peptides phosphorylated at low, middle, and higher concentrations yield low (green), medium (blue), and high (pink) affinity peptides selective for AKT1 (C) and AKT3 (D). Pathway analyses of the high-affinity peptides using EnrichR and the BioPlanet2019 database (E and F).

**Figure 5.**
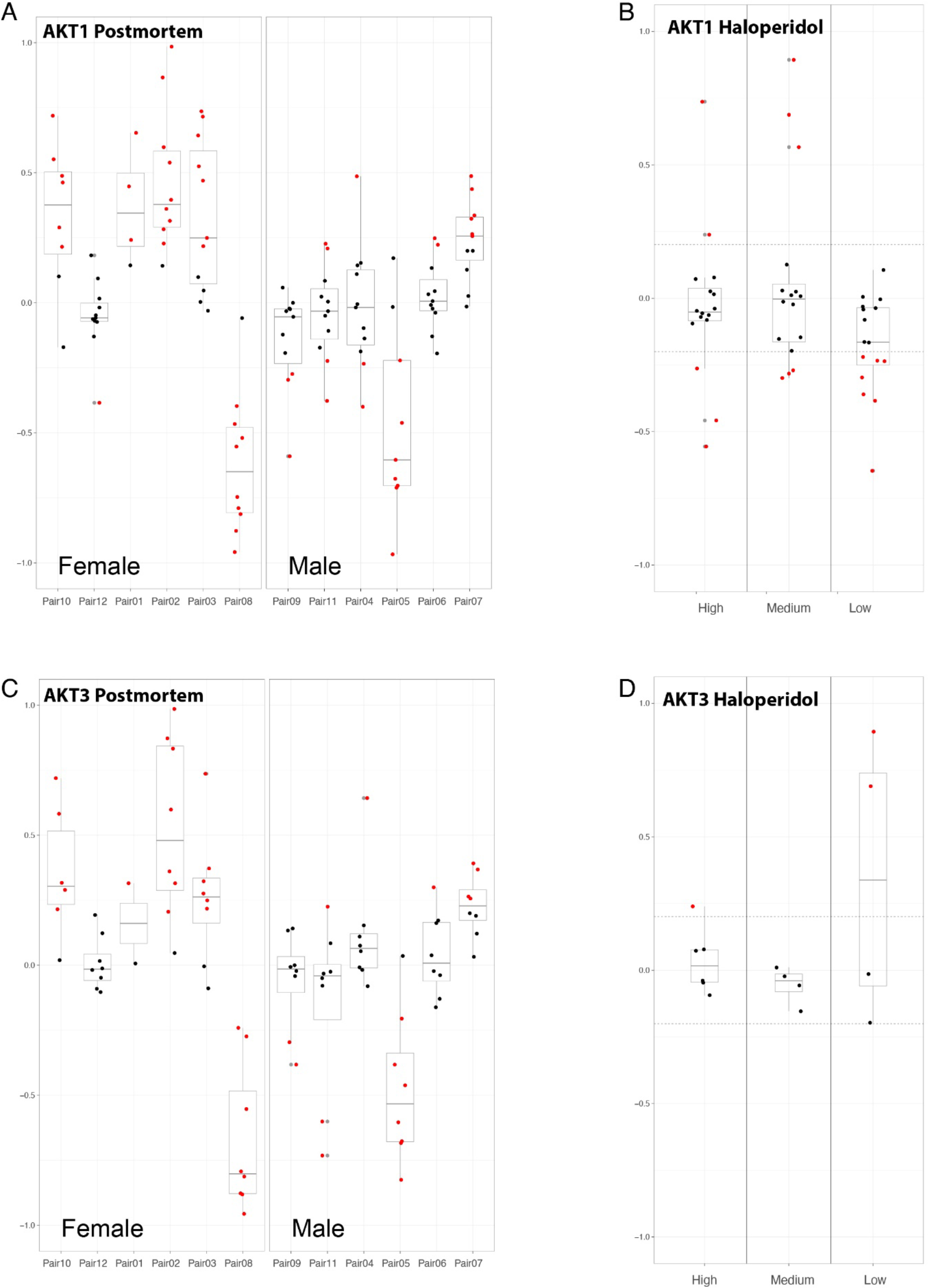
Reanalysis of published [23] PamGene Kinome array serine/threonine chip datasets from 12 pairs of schizophrenia and control subjects (anterior cingulate cortex tissue homogenate) and cortical homogenates from haloperidol treated rats datasets. Values for the recombinant AKT1 (A) and AKT3 (C) high-affinity peptides (Log2 fold change of case versus control) are represented as individual circles across the 12 subject pairs. For the rodent studies, Log2 fold change of case versus control for pooled samples (n = 10 per group) for low, medium, and high affinity peptides are shown (B, D). Red circles indicate increased or decreased activity levels greater than +/-15%.

### AKT1

Results of the clustering show that 17 peptides are classified as ‘high affinity’ (Fig 4A and S2). Principal component analysis (PCA) revealed that a majority of the differences between peptides in each cluster was attributed to the second dimension (PC2) with the first dimension (PC1) serving to separate clusters themselves; in this case, PC1 represented 94.3% of the variation (Fig 4C). 4/6 female subjects show higher activity (Fig 5A) for high-affinity AKT1 reporter peptides (pink circles, Fig 4C) in schizophrenia compared to age and gender-matched controls. Only one subject pair showed higher activity in the male group (Fig 5A). In addition, one female subject pair and one male subject pair show lower AKT1 activity in schizophrenia (Fig 5A). Finally, in the frontal cortex from haloperidol-decanoate treated rats (28.5 mg/kg for 9 months, q 3-week injections), phosphorylation of high-affinity AKT1 reporter peptides is equivocal, with 7 peptides unchanged, 2 increased, and 4 decreased (Fig 5B).

### AKT3

Initial results of the clustering show that 8 peptides are classified as ‘high affinity’ (Fig 4B and S2). Principal component analysis (PCA) revealed that a majority of the differences between peptides in each cluster was attributed to the second dimension (PC2) with the first dimension (PC1) serving to separate clusters themselves; in this case, PC1 represented 82.1% of the variation (Fig 4D). 3/6 female subjects show higher activity (Fig 5C) for high-affinity AKT3 reporter peptides (pink circles, Fig 4D) in schizophrenia compared to age and gender-matched controls. Only one subject pair showed higher activity in the male group (Fig 5C). In addition, one female subject pair and two male subject pairs show lower AKT3 activity in schizophrenia (Fig 5C). Finally, in the frontal cortex from haloperidol-decanoate treated rats (28.5 mg/kg for 9 months, q 3-week injections), phosphorylation of high-affinity AKT3 reporter peptides is equivocal, with 4 peptides unchanged, 1 increased, and zero decreased (Fig 5D).

### Pathway analyses

Notably, the STK array chip comprises 144 peptides, which map to 124 unique proteins, providing a small set of genes for pathway annotation. Despite this limitation, the analysis of pathways associated with AKT1 and AKT3 activity on the chip yields valuable insights. For AKT1, eight out of the top 10 pathways mapped to high-affinity peptides have been identified as dysregulated or involved in the pathogenesis of schizophrenia, including the activation of calcium-permeable kainate receptor, apoptotic DNA fragmentation and tissue homeostasis, and three pathways related to nitric oxide [40–42]. Additionally, three pathways are associated with ERBB2, a known upstream activator of AKT1 implicated in schizophrenia. These pathways are ERBB2 role in signal transduction and oncology, GRB7 events in ERBB2 signaling, and erythropoietin-mediated neuroprotection through NF-κB [43–45](Fig 4E). Similarly, for AKT3, nine out of the top 10 pathways mapped to high-affinity peptides are also implicated with schizophrenia, including regulation by CREM, GABA receptor activation, cell-extracellular matrix interactions, selenium metabolism, e-cadherin, ligand-gated ion channel transport, and three pathways related to AKT/mTOR/S6K signaling [46–52] (Fig 4-F).

Structurally, the high, medium and low-affinity AKT1 peptides show a large number of serine residues based on their sequence tags; a distinguishing factor between the low-affinity and other groups appears is an arginine at position 3. The high-affinity group also has an aspartate at position 9, with lysine at positions 8 and 10 (Fig S2A).

Structurally, the low, medium and high affinity AKT3 peptides show a large number of Serine residues as their defining factors. A distinguishing factor for the high affinity peptides is the RRR sequence present on positions 3, 4 and 5, while the low and medium affinity groups do not have R at position 5 (Fig S2B).

## Discussion

While AKT1-3 transcripts have markedly differential cellular expression patterns (Supplementary material, Tables 2-5), we found increased mRNA expression for all three AKT isotypes in neurons in schizophrenia. This result was unexpected as prior animal studies suggest specific functions for the different isotypes [14, 15, 53, 54]. One possibility is that isoform specific regulation of AKT1-3 does not occur at the level of transcription. Alternatively, our findings may reflect a global deficit or compensation for impaired insulin signaling, a state believed to be prevalent in schizophrenia [13, 21, 55]. Given the prominence of insulin signaling pathways in biological systems, changes in AKT mRNA gene expression may reflect a compensatory response to perturbations in this critical and tightly regulated cellular process.

Changes in AKT gene expression appear to be relatively sex-specific, with changes in female samples accounting for most of the effect (Figs 5 and S1). Consistent with this observation, we found higher kinase activity for AKT1 reporter peptides in 4 out of 6 female subject pairs, but only 1 out of 6 matched male pairs (Fig 5). We observed a similar pattern for AKT3 (Fig 5). These results suggest divergent roles for AKT in female versus male subjects with schizophrenia.

Examination of previously published RNAseq and microarray datasets revealed few alterations in neuronal expression of the AKT transcripts in schizophrenia. Congruent with our findings, one study in the superior temporal gyrus (STG) found increased neuronal AKT1 mRNA expression [56]. In contrast, one ACC and three DLPFC studies found no changes in neuron-specific transcript expression for any of the AKT isotypes [34, 57, 58]. These divergent findings may be secondary to differences in subject demographics, brain region, treatment with psychotropic medications, and/or substance use history [24, 25].

It is widely accepted that metabolic dysfunction plays a key role in the etiopathogenesis of schizophrenia [13, 21, 55, 59]. Alterations in glucose utilization, secondary to perturbed insulin signaling pathways, likely contributes to the cognitive deficits seen in schizophrenia [12, 60, 61]. AKT activity is dependent on activation of upstream growth factor (such as insulin) receptor occupancy and activation [21, 62, 63]. Once active, AKT may then regulate glucose metabolism via phosphorylation-mediated inhibition of the constitutively active GSK3-B, which promotes glycogen and protein synthesis [21, 62, 63]. Since insulin signaling regulates AKT activity, a primary deficit of insulin signaling pathways might lead to increased AKT mRNA expression as a compensatory response. This leads to the question of whether AKT protein expression reflects changes in mRNA levels. We previously measured region level AKT protein expression (using a pan-specific antibody) and did not detect changes in total protein expression in the ACC [23]. Several other studies found no changes in total protein expression in postmortem prefrontal cortex [23, 47, 64–67] while five others found decreased total AKT protein [5, 16, 18, 19, 21].

Since AKT is an enzyme, measuring expression levels may not be the best way to assess changes in activity. Previously, we and others have found region-level decreases in phospho-AKT expression, a proxy for AKT activity, in the ACC and DLPFC in schizophrenia [19, 21, 23, 53], while another found no differences in the PFC [47]. Taken together, these findings suggest there may be region specific decreases in AKT protein expression in schizophrenia, with a decrease in phospho-AKT levels suggesting lower enzyme activity.

Notably, these previously published AKT protein studies were not cell or isoform-specific, limiting the interpretation of these findings. If these previous region-level findings for AKT protein expression extend to neurons, the increases in neuronal AKT1-3 mRNAs found in the present study support a hypothesis of pathological uncoupling of AKT transcript and protein expression. Given the perturbation of insulin signaling in schizophrenia [55, 68, 69], neurons may be transcribing more mRNA in an attempt to compensate for decreased AKT activity and/or downstream effects of lower AKT signaling including diminished glucose utilization [70, 71].

While transcript, protein, and phospho-protein expression do not always correlate [72, 73], protein function is the ultimate biological effect of an enzyme. Previously, in region-level ACC tissue homogenates, we found increased AKT-specific activity, with no change in total AKT kinase activity [23]. Current approaches make an efficient assessment of kinase activity at the cell level infeasible (time and expense) in the postmortem brain, particularly for AKT isoforms. We did however perform experiments to inform region-level activity for the AKT1 and AKT3 isoforms using a recombinant kinase approach. We assayed recombinant AKT1 and AKT3 activity levels on a peptide array to empirically determine the peptides that best report recombinant AKT1 and AKT3 activity (Fig 4). Next, we interrogated a previously published [23] postmortem study using the same brain region from the same brain bank, albeit a different cohort of subjects (Fig 5). Surprisingly, we found evidence of higher AKT1 activity in 4/6 female subject pairs and 1/6 male subject pairs, with lower activity patterns in one female and one male subject pair. A similar pattern was observed for AKT3.

These data suggest that global AKT1 and AKT3 activity may be increased in some subjects with schizophrenia. Given we previously found no changes in AKT total activity (and a decrease in pan phospho-AKT levels), there may be differential changes in AKT1, versus AKT2 and/or AKT3 kinase activity. This conjecture is supported by the differential expression of AKT1-3 mRNAs in neurons and other cell types in normal brain, with AKT3 > AKT2 > AKT1 in neurons (https://www.brainrnaseq.org/)(Supplementary Tables 2-5). We were not able to assess AKT2 with the recombinant profile approach as recombinant AKT2 did not give sufficient signal on the peptide array. Preclinical studies suggest diverse cellular expressions and functions for AKT1-3, further supporting this hypothesis [3]. Assessment of cell-subtype specific kinase activity in schizophrenia is an important next step to determine the relative contributions of AKT1-3 to perturbations of signaling networks in schizophrenia.

Since AKT is a hub in complex signaling networks, investigation of its pathway components may provide additional insight into the pathophysiology of schizophrenia. One such component is 3-phosphoinositide-dependent protein kinase (PDPK1 aka PDK1), an upstream activator of AKT [74]. We found increased neuronal PDK1 mRNA expression in schizophrenia, consistent with the hypothesis that there is a compensatory response driving the upregulation of genes associated with the insulin signaling pathway.

To complement our findings for protein kinases, we also studied several protein phosphatases known to regulate AKT, including PPP2CA, PHLPP1, PHLPP2, and PTEN [75–80]. Interestingly, we found increased expression of mRNAs for PHLPP2 and PTEN in pyramidal neurons and ACC homogenate and increased expression of PPP2CA only in the ACC homogenate. These findings suggest a complexity beyond a straightforward balance between kinase versus phosphatase activity; our study lacks resolution in terms of assessing the subcellular colocalization of these gene products, or even if these changes in cellular transcripts reflect changes in protein expression or enzyme activity. Regardless of the precise mechanism, there appears to be a potent compensatory response driving increased expression of transcripts for insulin signaling pathway genes in pyramidal neurons in chronic schizophrenia.

We also found increased expression of transcripts for Forkhead box protein O1 (FOXO1) in pyramidal neurons and ACC homogenate. FOXO1 is a transcription factor that regulates gluconeogenesis and glycogenolysis via insulin signaling pathways, downstream of AKT and other kinases and phosphatases in this canonical signaling cascade [81, 82]. Elevated expression of FOXO1 may be driving transcript expression of AKT and related genes in neurons in schizophrenia, supporting the hypothesis of insulin signaling dysregulation and a compensatory response at the level of transcript expression.

To assess the impact of antipsychotic medications on our dependent measures, we deployed a “look up” approach, using >50 antipsychotic treatment signatures, covering myriad doses and types of antipsychotics. Among our nine genes of interest, we found that typical and atypical antipsychotics had minimal effects on mRNA expression in the frontal cortex. One dataset had decreased FOXO1 mRNA expression with an atypical antipsychotic in the frontal cortex [83]. However, we found increased FOXO1 expression in schizophrenia subjects, suggesting chronic antipsychotic treatment did not drive this result. For the remainder of the genes of interest, antipsychotics (typical or atypical) had minimal effects on transcript expression, suggesting our transcript findings in schizophrenia subjects were not influenced by antipsychotic treatment.

Previous studies have investigated the effects of antipsychotic treatment on AKT expression. One study treated rats with varying dosages of haloperidol (0.5, 1, 2, or 4 mg/kg), clozapine (5, 10, or 20 mg/kg), or vehicle (0.3% tartaric acid in saline) and found a significant increase in phospho-AKT expression in the frontal cortex for all doses at acute 15 and 30-minute timepoints; notably, expression levels returned to baseline after one hour. In contrast, in the same study rats treated with clozapine showed a significant increase in frontal cortex phospho-AKT protein expression at all time points and doses [84]. Studies with longer duration of antipsychotic treatment have found different results. Rats treated twice daily for 21 days with haloperidol (0.5 mg/kg), clozapine (5 mg/kg), or vehicle (1 ml/kg) had no changes in total AKT protein expression but decreased phospho-AKT in the frontal cortex [47]. Finally, we found no changes in phospho-AKT in the frontal cortex brain homogenate following nine months of treatment with haloperidol [23]. These data suggest that while acute antipsychotic treatment may impact region-level phospho-AKT levels, these changes appear to normalize with chronic treatment. In chronic schizophrenia, subjects are often taking antipsychotic medications for decades, most analogous to the aforementioned chronic treatment studies in rats [23].

We previously reported that frontal cortical homogenate from rats treated with haloperidol for 9 months had decreased AKT-specific activity, with no change in total activity [23]. To our knowledge, AKT activity assays directly assessing the effects of atypical antipsychotics have not been performed. Taken together, the present data and previously published findings support the conclusion that increases in AKT mRNAs in neurons and changes in AKT activity in ACC homogenate are likely not secondary to antipsychotic treatment.

While translationally valuable, the use of postmortem tissue has limitations. Postmortem cohorts are typically well-matched for age, sex, PMI, pH, and RIN but are relatively heterogenous compared to animal models. We explored cell-level changes in transcript expression, which does not always predict changes in protein expression or activity [72, 73]. We used LMD to provide cell subtype specificity; this technique relies on morphological identification of cells and yields a pooled sample of cells with a pyramidal neuron profile. Other more sophisticated approaches are available, including single nuclei RNAseq [85]. We deployed LMD-QPCR since we sought to test a specific hypothesis across a population of cells, rather than examining profiles of subsets/subtypes of pyramidal neurons. Further, protein studies using LMD are prohibitive as it takes about 10,000 captured cells per subject to get a measurable band on a Western blot or about 1,000 captured cells per subject to perform standard biochemical kinase activity assays. While our recombinant AKT studies provide an important tool for assessing changes in AKT1 and AKT3 activity on the peptide activity array, we were unable to generate similar signatures with available recombinant AKT2 on this same platform.

In summary, we postulate that schizophrenia occurs due to an accumulation of many subtle changes in genes and signaling networks that interfere with the function of crucial biological processes. Thus, understanding the dynamics of AKT and related signaling pathways in the pathophysiology of schizophrenia is essential for the identification of novel targets for therapeutics. Currently, there are no FDA approved (or in clinical trials) AKT-targeting drugs for schizophrenia. However AKT inhibitors are being developed as approaches to treat cancer [86], and the medicinal flavanone glycoside naringin may exert its protective effects via alteration of wnt/β-catenin and Akt/GSK-3 β pathways [87]. In conclusion, our findings suggest that persons with chronic schizophrenia have increases in neurons in transcript expression of insulin signaling pathway genes, including protein kinases, transcription factors, and phosphatases. We suspect these changes are a compensatory response to a primary defect of insulin signaling (or insulin resistance), leading cells to attempt to increase gene expression of hub proteins for these pathways.

## Acknowledgements

This work was supported by the National Institute of Mental Health (MH107487, MH121102 to REM) and National Institute of Aging (AG057598 to REM)

## Notes

### Competing Interest Statement

The authors have declared no competing interest.

## References

1. Staal, S.P., Molecular cloning of the akt oncogene and its human homologues AKT1 and AKT2: amplification of AKT1 in a primary human gastric adenocarcinoma. Proc Natl Acad Sci U S A, 1987. 84(14): p. 5034–7.

2. Cohen, M.M., Jr., The AKT genes and their roles in various disorders. Am J Med Genet A, 2013. 161a(12): p. 2931-7.

3. Dummler, B., et al., Life with a single isoform of Akt: mice lacking Akt2 and Akt3 are viable but display impaired glucose homeostasis and growth deficiencies. Mol Cell Biol, 2006. 26(21): p. 8042–51.

4. Arguello, P.A. and J.A. Gogos, A signaling pathway AKTing up in schizophrenia. J Clin Invest, 2008. 118(6): p. 2018–21.

5. Emamian, E.S., et al., Convergent evidence for impaired AKT1-GSK3beta signaling in schizophrenia. Nat Genet, 2004. 36(2): p. 131–7.

6. Emamian, E.S., AKT/GSK3 signaling pathway and schizophrenia. Front Mol Neurosci, 2012. 5: p. 33.

7. Green, M.F. and P.D. Harvey, Cognition in schizophrenia: Past, present, and future.

8. Schizophr Res Cogn, 2014. 1(1): p. e1–e9.

9. Nuechterlein, K.H., et al., Identification of separable cognitive factors in schizophrenia. Schizophr Res, 2004. 72(1): p. 29–39.

10. Lai, W.S., et al., Akt1 deficiency affects neuronal morphology and predisposes to abnormalities in prefrontal cortex functioning. Proc Natl Acad Sci U S A, 2006. 103(45): p. 16906–11.

11. Lin, C.H., et al., A role for the PI-3 kinase signaling pathway in fear conditioning and synaptic plasticity in the amygdala. Neuron, 2001. 31(5): p. 841–51.

12. Cho, H., et al., Insulin resistance and a diabetes mellitus-like syndrome in mice lacking the protein kinase Akt2 (PKB beta). Science, 2001. 292(5522): p. 1728-31.

13. Wijtenburg, S.A., et al., Brain insulin resistance and altered brain glucose are related to memory impairments in schizophrenia. Schizophr Res, 2019. 208: p. 324–330.

14. Sullivan, C.R., et al., Defects in Bioenergetic Coupling in Schizophrenia. Biol Psychiatry, 2018. 83(9): p. 739–750.

15. Howell, K.R., K. Floyd, and A.J. Law, PKBgamma/AKT3 loss-of-function causes learning and memory deficits and deregulation of AKT/mTORC2 signaling: Relevance for schizophrenia. PLoS One, 2017. 12(5): p. e0175993.

16. Bergeron, Y., et al., Genetic Deletion of Akt3 Induces an Endophenotype Reminiscent of Psychiatric Manifestations in Mice. Front Mol Neurosci, 2017. 10: p. 102.

17. Ikeda, M., et al., Association of AKT1 with schizophrenia confirmed in a Japanese population. Biol Psychiatry, 2004. 56(9): p. 698–700.

18. Biological insights from 108 schizophrenia-associated genetic loci. Nature, 2014. 511(7510): p. 421-7.

19. Thiselton, D.L., et al., AKT1 is associated with schizophrenia across multiple symptom dimensions in the Irish study of high density schizophrenia families. Biol Psychiatry, 2008. 63(5): p. 449–57.

20. Chadha, R. and J.H. Meador-Woodruff, Downregulated AKT-mTOR signaling pathway proteins in dorsolateral prefrontal cortex in Schizophrenia. Neuropsychopharmacology, 2020. 45(6): p. 1059–1067.

21. Kunii, Y., et al., Evidence for Altered Phosphoinositide Signaling-Associated Molecules in the Postmortem Prefrontal Cortex of Patients with Schizophrenia. Int J Mol Sci, 2021. 22(15).

22. Zhao, Z., et al., Insulin receptor deficits in schizophrenia and in cellular and animal models of insulin receptor dysfunction. Schizophr Res, 2006. 84(1): p. 1–14.

23. McGuire, J.L., et al., Altered serine/threonine kinase activity in schizophrenia. Brain Res, 2014. 1568: p. 42–54.

24. McGuire, J.L., et al., Abnormalities of signal transduction networks in chronic schizophrenia. NPJ Schizophr, 2017. 3(1): p. 30.

25. McCullumsmith, R.E., et al., Postmortem brain: an underutilized substrate for studying severe mental illness. Neuropsychopharmacology, 2014. 39(1): p. 65–87.

26. McCullumsmith, R.E. and J.H. Meador-Woodruff, Novel approaches to the study of postmortem brain in psychiatric illness: old limitations and new challenges. Biol Psychiatry, 2011. 69(2): p. 127–33.

27. Boland, E., et al., Mapping of deletion and translocation breakpoints in 1q44 implicates the serine/threonine kinase AKT3 in postnatal microcephaly and agenesis of the corpus callosum. Am J Hum Genet, 2007. 81(2): p. 292–303.

28. Tan, H.Y., et al., Genetic variation in AKT1 is linked to dopamine-associated prefrontal cortical structure and function in humans. J Clin Invest, 2008. 118(6): p. 2200–8.

29. Lin, J.R., et al., Integrated Post-GWAS Analysis Sheds New Light on the Disease Mechanisms of Schizophrenia. Genetics, 2016. 204(4): p. 1587–1600.

30. Trubetskoy, V., et al., Mapping genomic loci implicates genes and synaptic biology in schizophrenia. Nature, 2022. 604(7906): p. 502-508.

31. McCullumsmith, R.E., et al., Cell-specific abnormalities of glutamate transporters in schizophrenia: sick astrocytes and compensating relay neurons? Mol Psychiatry, 2016. 21(6): p. 823–30.

32. Sodhi, M.S., et al., Glutamatergic gene expression is specifically reduced in thalamocortical projecting relay neurons in schizophrenia. Biol Psychiatry, 2011. 70(7): p. 646–54.

33. O’Donovan, S.M., et al., Cell-subtype-specific changes in adenosine pathways in schizophrenia. Neuropsychopharmacology, 2018. 43(8): p. 1667–1674.

34. Sullivan, C.R., et al., Neuron-specific deficits of bioenergetic processes in the dorsolateral prefrontal cortex in schizophrenia. Mol Psychiatry, 2019. 24(9): p. 1319–1328.

35. Wu, X., et al., Transcriptional profile of pyramidal neurons in chronic schizophrenia reveals lamina-specific dysfunction of neuronal immunity. Mol Psychiatry, 2021. 26(12): p. 7699–7708.

36. Asah, S., et al., A bioinformatic inquiry of the EAAT2 interactome in postmortem and neuropsychiatric datasets. Schizophr Res, 2020.

37. Castellani, L.N., et al., Antipsychotics impair regulation of glucose metabolism by central glucose. Mol Psychiatry, 2022. 27(11): p. 4741–4753.

38. Creeden, J.F., et al., Kinome Array Profiling of Patient-Derived Pancreatic Ductal Adenocarcinoma Identifies Differentially Active Protein Tyrosine Kinases. Int J Mol Sci, 2020. 21(22).

39. DePasquale, E.A.K., et al., KRSA: An R package and R Shiny web application for an end-to-end upstream kinase analysis of kinome array data. PLoS One, 2021. 16(12): p. e0260440.

40. Schrode, N., et al., Synergistic effects of common schizophrenia risk variants. Nat Genet, 2019. 51(10): p. 1475–1485.

41. Meador-Woodruff, J.H., K.L. Davis, and V. Haroutunian, Abnormal kainate receptor expression in prefrontal cortex in schizophrenia. Neuropsychopharmacology, 2001. 24(5): p. 545–52.

42. Nasyrova, R.F., et al., Role of nitric oxide and related molecules in schizophrenia pathogenesis: biochemical, genetic and clinical aspects. Front Physiol, 2015. 6: p. 139.

43. Notaras, M., et al., Schizophrenia is defined by cell-specific neuropathology and multiple neurodevelopmental mechanisms in patient-derived cerebral organoids. Mol Psychiatry, 2022. 27(3): p. 1416–1434.

44. Harrison, P.J. and D.R. Weinberger, Schizophrenia genes, gene expression, and neuropathology: on the matter of their convergence. Mol Psychiatry, 2005. 10(1): p. 40–68; image 5.

45. Murphy, C.E., A.K. Walker, and C.S. Weickert, Neuroinflammation in schizophrenia: the role of nuclear factor kappa B. Transl Psychiatry, 2021. 11(1): p. 528.

46. Seshadri, S., et al., Disrupted-in-Schizophrenia-1 expression is regulated by beta-site amyloid precursor protein cleaving enzyme-1-neuregulin cascade. Proc Natl Acad Sci U S A, 2010. 107(12): p. 5622–7.

47. Forero, D.A., et al., A network of synaptic genes associated with schizophrenia and bipolar disorder. Schizophr Res, 2016. 172(1-3): p. 68–74.

48. Ibarra-Lecue, I., et al., Ribosomal Protein S6 Hypofunction in Postmortem Human Brain Links mTORC1-Dependent Signaling and Schizophrenia. Front Pharmacol, 2020. 11: p. 344.

49. de Jonge, J.C., et al., GABAergic Mechanisms in Schizophrenia: Linking Postmortem and In Vivo Studies. Front Psychiatry, 2017. 8: p. 118.

50. Pantazopoulos, H., et al., Molecular signature of extracellular matrix pathology in schizophrenia. Eur J Neurosci, 2021. 53(12): p. 3960–3987.

51. Maes, M., et al., First Episode Psychosis and Schizophrenia Are Systemic Neuro-Immune Disorders Triggered by a Biotic Stimulus in Individuals with Reduced Immune Regulation and Neuroprotection. Cells, 2021. 10(11).

52. Imbrici, P., D.C. Camerino, and D. Tricarico, Major channels involved in neuropsychiatric disorders and therapeutic perspectives. Front Genet, 2013. 4: p. 76.

53. Pitts, M.W., A.V. Raman, and M.J. Berry,Schizophrenia, Oxidative Stress and Selenium, in Selenium: Its Molecular Biology and Role in Human Health, D.L. Hatfield, M.J. Berry, and V.N. Gladyshev, Editors. 2012, Springer New York: New York, NY. p. 355-367.

54. Balu, D.T., et al., Akt1 deficiency in schizophrenia and impairment of hippocampal plasticity and function. Hippocampus, 2012. 22(2): p. 230–40.

55. Leibrock, C., et al., Akt2 deficiency is associated with anxiety and depressive behavior in mice. Cell Physiol Biochem, 2013. 32(3): p. 766–77.

56. Henkel, N.D., et al., Schizophrenia: a disorder of broken brain bioenergetics. Mol Psychiatry, 2022. 27(5): p. 2393–2404.

57. Pietersen, C.Y., et al., Molecular profiles of pyramidal neurons in the superior temporal cortex in schizophrenia. J Neurogenet, 2014. 28(1-2): p. 53–69.

58. Arion, D., et al., Distinctive transcriptome alterations of prefrontal pyramidal neurons in schizophrenia and schizoaffective disorder. Mol Psychiatry, 2015. 20(11): p. 1397–1405.

59. Arion, D., et al., Transcriptome Alterations in Prefrontal Pyramidal Cells Distinguish Schizophrenia From Bipolar and Major Depressive Disorders. Biol Psychiatry, 2017. 82(8): p. 594–600.

60. Bryll, A., et al., Oxidative-Antioxidant Imbalance and Impaired Glucose Metabolism in Schizophrenia. Biomolecules, 2020. 10(3).

61. Zhang, X., et al., Glucose disturbances, cognitive deficits and white matter abnormalities in first-episode drug-naive schizophrenia. Mol Psychiatry, 2020. 25(12): p. 3220–3230.

62. Tang, S.X., et al., Metabolic disturbances, hemoglobin A1c, and social cognition impairment in Schizophrenia spectrum disorders. Transl Psychiatry, 2022. 12(1): p. 233.

63. Sharma, M. and C.S. Dey, Role of Akt isoforms in neuronal insulin signaling and resistance. Cell Mol Life Sci, 2021. 78(23): p. 7873–7898.

64. Zheng, W., et al., The possible role of the Akt signaling pathway in schizophrenia. Brain Res, 2012. 1470: p. 145–58.

65. Karege, F., et al., Association of AKT1 gene variants and protein expression in both schizophrenia and bipolar disorder. Genes Brain Behav, 2010. 9(5): p. 503–11.

66. Hino, M., et al., Decreased VEGFR2 expression and increased phosphorylated Akt1 in the prefrontal cortex of individuals with schizophrenia. J Psychiatr Res, 2016. 82: p. 100–8.

67. Ide, M., et al., Failure to support a genetic contribution of AKT1 polymorphisms and altered AKT signaling in schizophrenia. J Neurochem, 2006. 99(1): p. 277–87.

68. Amar, S., et al., Possible involvement of post-dopamine D2 receptor signalling components in the pathophysiology of schizophrenia. Int J Neuropsychopharmacol, 2008. 11(2): p. 197–205.

69. Agarwal, S.M., et al., Brain insulin action: Implications for the treatment of schizophrenia. Neuropharmacology, 2020. 168: p. 107655.

70. Ding, H., et al., Shared genetics of psychiatric disorders and type 2 diabetes:a large-scale genome-wide cross-trait analysis. J Psychiatr Res, 2023. 159: p. 185–195.

71. Mackenzie, R.W. and B.T. Elliott, Akt/PKB activation and insulin signaling: a novel insulin signaling pathway in the treatment of type 2 diabetes. Diabetes Metab Syndr Obes, 2014. 7: p. 55–64.

72. Zhang, Z., H. Liu, and J. Liu, Akt activation: A potential strategy to ameliorate insulin resistance. Diabetes Res Clin Pract, 2019. 156: p. 107092.

73. de Sousa Abreu, R., et al., Global signatures of protein and mRNA expression levels. Mol Biosyst, 2009. 5(12): p. 1512–26.

74. Vogel, C. and E.M. Marcotte, Insights into the regulation of protein abundance from proteomic and transcriptomic analyses. Nat Rev Genet, 2012. 13(4): p. 227–32.

75. Brunet, A., S.R. Datta, and M.E. Greenberg, Transcription-dependent and-independent control of neuronal survival by the PI3K-Akt signaling pathway. Curr Opin Neurobiol, 2001. 11(3): p. 297–305.

76. Andreozzi, F., et al., Increased levels of the Akt-specific phosphatase PH domain leucine-rich repeat protein phosphatase (PHLPP)-1 in obese participants are associated with insulin resistance. Diabetologia, 2011. 54(7): p. 1879–87.

77. Zhao, L., R. Li, and Y.H. Gan, Knockdown of Yin Yang 1 enhances anticancer effects of cisplatin through protein phosphatase 2A-mediated T308 dephosphorylation of AKT. Cell Death Dis, 2018. 9(7): p. 747.

78. Zeng, Q., et al., The miR-345-3p/PPP2CA signaling axis promotes proliferation and invasion of breast cancer cells. Carcinogenesis, 2022. 43(2): p. 150–159.

79. Liao, Y. and M.C. Hung, A new role of protein phosphatase 2a in adenoviral E1A protein-mediated sensitization to anticancer drug-induced apoptosis in human breast cancer cells. Cancer Res, 2004. 64(17): p. 5938–42.

80. Nowak, D.G., et al., The PHLPP2 phosphatase is a druggable driver of prostate cancer progression. J Cell Biol, 2019. 218(6): p. 1943–1957.

81. Bu, L., et al., PTEN suppresses tumorigenesis by directly dephosphorylating Akt. Signal Transduct Target Ther, 2021. 6(1): p. 262.

82. Thiel, G., L.A. Guethlein, and O.G. Rossler, Insulin-Responsive Transcription Factors. Biomolecules, 2021. 11(12).

83. Kousteni, S., FoxO1, the transcriptional chief of staff of energy metabolism. Bone, 2012. 50(2): p. 437–43.

84. C, C. Ventral hippocampal lesion, tetrodotoxin disruption of the ventral hippocampus, and chronic administration of neuroleptics. Jan 13, 2006; Available from: https://www.ncbi.nlm.nih.gov/geo/query/acc.cgi?acc=GSE4031.

85. Roh, M.S., et al., Haloperidol and clozapine differentially regulate signals upstream of glycogen synthase kinase 3 in the rat frontal cortex. Exp Mol Med, 2007. 39(3): p. 353–60.

86. Shukla, R., et al., Signature-based approaches for informed drug repurposing: targeting CNS disorders. Neuropsychopharmacology, 2021. 46(1): p. 116–130.

87. Howell, S.J., et al., Fulvestrant plus capivasertib versus placebo after relapse or progression on an aromatase inhibitor in metastatic, oestrogen receptor-positive, HER2-negative breast cancer (FAKTION): overall survival, updated progression-free survival, and expanded biomarker analysis from a randomised, phase 2 trial. The Lancet Oncology, 2022. 23(7): p. 851-864.

88. George, M.Y., et al., Potential therapeutic antipsychotic effects of Naringin against ketamine-induced deficits in rats: involvement of Akt/GSK-3β and Wnt/β-catenin signaling pathways. Life sciences, 2020. 249: p. 117535.

