## Supplemental Tables and Figures for "Neuronal alterations in AKT isotype expression in schizophrenia"

### Supplemental Information

#### Supplementary Table 1. Human Taqman primers

| **Assay** | **Gene Name** | **Primer** |
| --- | --- | --- |
| AKT serine/threonine kinase 1 | AKT1 | Hs00178289_m1 |
| AKT serine/threonine kinase 2 | AKT2 | Hs01086102_m1 |
| AKT serine/threonine kinase 3 | AKT3 | Hs00987350_m1 |
| Forkhead box O1 | FOXO1 | Hs00231106_m1 |
| Phosphatase and tensin homolog | PTEN | Hs02621230_s1 |
| 3-phosphoinositide dependent protein kinase 1 | PDPK1 | Hs00928927_m1 |
| PH domain and leucine rich repeat protein phosphatase 1 | PHLPP1 | Hs01597875_m1 |
| PH domain and leucine rich repeat protein phosphatase 2 | PHLPP2 | Hs00982295_m1 |
| Protein phosphatase 2 catalytic subunit alpha | PPP2CA | Hs00427260_m1 |
| Cyclophillin A | PPIA | Hs99999904_m1 |
| Beta-actin | ACTB | Hs99999903_m1 |
| Beta-2-microglobulin | B2M | Hs99999907_m1 |
| Glyceraldehyde-3-phosphate dehydrogenase | GAPDH | Hs99999905_m1 |

List of Taqman primers used to measure gene expression targets in postmortem tissue.

#### Supplementary Table 2. BRAINseq data for AKT1-3

| HGNC_Symbol | CellType | log10(FPKM+1) | Species |
|---|---|---|---|
| AKT1 | Astrocytes | 1.48 | Mouse |
| AKT1 | Endothelial Cells | 1.82 | Mouse |
| AKT1 | Microglia | 2.04 | Mouse |
| AKT1 | Myelinating Oligodendrocytes | 1.35 | Mouse |
| AKT1 | Neuron | 1.75 | Mouse |
| AKT1 | Newly Formed Oligodendrocyte | 1.62 | Mouse |
| AKT1 | Oligodendrocyte Precursor Cell | 1.77 | Mouse |
| AKT2 | Astrocytes | 1.64 | Mouse |
| AKT2 | Endothelial Cells | 0.67 | Mouse |
| AKT2 | Microglia | 1.26 | Mouse |
| AKT2 | Myelinating Oligodendrocytes | 0.92 | Mouse |
| AKT2 | Neuron | 0.99 | Mouse |
| AKT2 | Newly Formed Oligodendrocyte | 1.03 | Mouse |
| AKT2 | Oligodendrocyte Precursor Cell | 1.24 | Mouse |
| AKT3 | Astrocytes | 1.18 | Mouse |
| AKT3 | Endothelial Cells | 1.31 | Mouse |
| AKT3 | Microglia | 0.44 | Mouse |
| AKT3 | Myelinating Oligodendrocytes | 0.8 | Mouse |
| AKT3 | Neuron | 1.51 | Mouse |
| AKT3 | Newly Formed Oligodendrocyte | 1 | Mouse |
| AKT3 | Oligodendrocyte Precursor Cell | 1.19 | Mouse |
| AKT1 | Endothelial | 0.45 | Human |
| AKT1 | FetalAstrocytes | 0.77 | Human |
| AKT1 | MatureAstrocytes | 0.26 | Human |
| AKT1 | Microglia_Macrophage | 0.18 | Human |
| AKT1 | Neurons | 0.11 | Human |
| AKT1 | Oligodendrocytes | 0.19 | Human |
| AKT2 | Endothelial | 0.22 | Human |
| AKT2 | FetalAstrocytes | 0.86 | Human |
| AKT2 | MatureAstrocytes | 0.47 | Human |
| AKT2 | Microglia_Macrophage | 0.16 | Human |
| AKT2 | Neurons | 0.18 | Human |
| AKT2 | Oligodendrocytes | 0.25 | Human |
| AKT3 | Endothelial | 1.81 | Human |
| AKT3 | FetalAstrocytes | 1.45 | Human |
| AKT3 | MatureAstrocytes | 1.37 | Human |
| AKT3 | Microglia_Macrophage | 0.99 | Human |
| AKT3 | Neurons | 1.67 | Human |
| AKT3 | Oligodendrocytes | 1.68 | Human |

**Supplementary Table 3**. **AKT1 Brain RNAseq Data**

| AKT 1 | |
|---|---|
| **Cell Type** | **log10 (FPKM+1)** |
| Endothelial | 0.45 |
| Fetal Astrocytes | 0.77 |
| Mature Astrocytes | 0.26 |
| Microglia Macrophage | 0.18 |
| Neurons | 0.11 |
| Oligodendrocytes | 0.19 |

Gene expression values log10 (FPKM+1) per cell type in human brain.

**Supplementary Table 4. AKT2 Brain RNAseq Data**

| AKT 2 | |
|---|---|
| **Cell Type** | **log10 (FPKM+1)** |
| Endothelial | 0.22 |
| Fetal Astrocytes | 0.86 |
| Mature Astrocytes | 0.47 |
| Microglia Macrophage | 0.16 |
| Neurons | 0.18 |
| Oligodendrocytes | 0.25 |

Gene expression values log10 (FPKM+1) per cell type in human brain.

**Supplementary Table 5**. **AKT3 Brain RNAseq Data**

| AKT 3 | |
|---|---|
| **Cell Type** | **log10 (FPKM+1)** |
| Endothelial | 1.81 |
| Fetal Astrocytes | 1.45 |
| Mature Astrocytes | 1.37 |
| Microglia Macrophage | 0.99 |
| Neurons | 1.67 |
| Oligodendrocytes | 1.68 |

Gene expression values log10 (FPKM+1) per cell type in human brain.

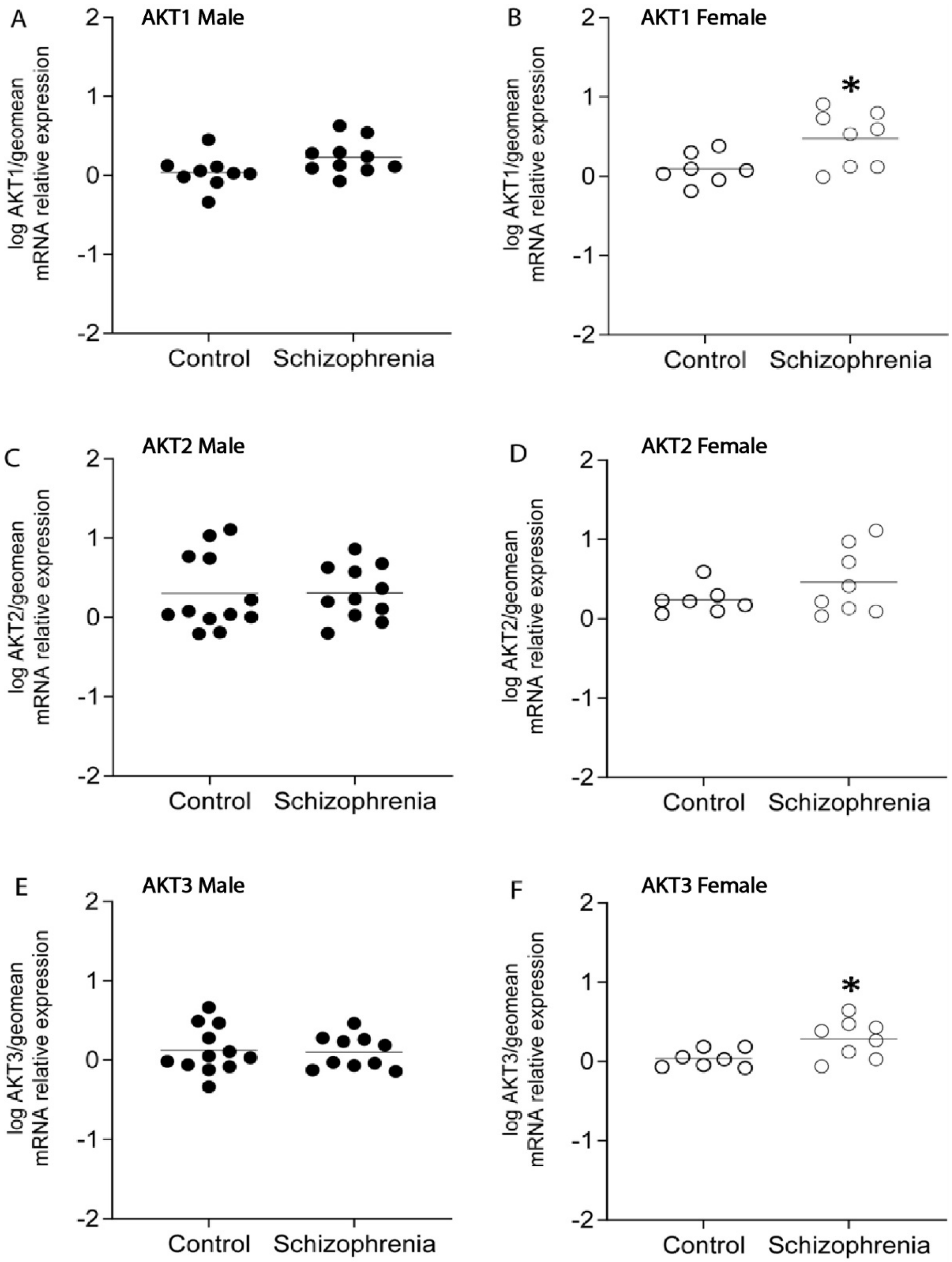


**Supplementary Figure 1. Enriched pyramidal cell population expression of AKT serine/threonine protein kinase isotypes in males and females**. Open circles indicate females, closed circles indicate males. Analysis revealed increased expression of B) AKT1 female and F) AKT3 female schizophrenia subjects compared to controls (*$p<0.05$). There was no significant change in gene expression of A) AKT1 male, C) AKT2 male, D) AKT2 female, or E) AKT3 male schizophrenia subjects compared to controls. Data are log transformed and analyzed using either Student's t-test or Mann-Whitney test. Data mean+/- SEM, n=7-12/group. (AKT1: AKT serine/threonine kinase 1, AKT2: AKT serine/threonine kinase 2, AKT3: AKT serine/threonine kinase 3)

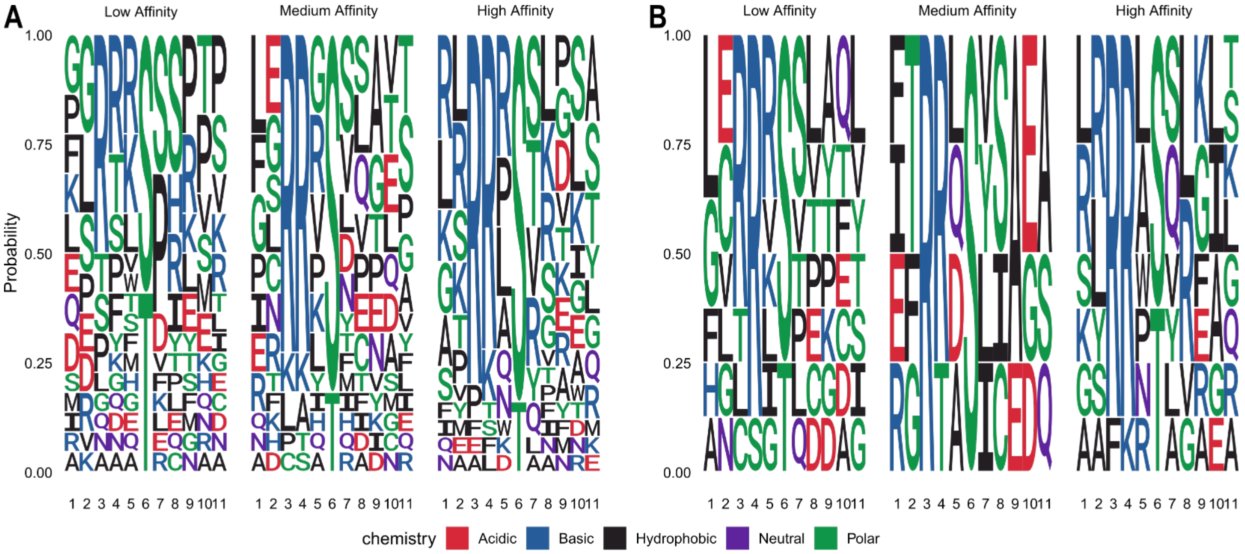


C

| Peptide | Sequence | Name | Symbol |
|---|---|---|---|
| RS6_228_240 | IAKRRRLSSLRAS | 40S ribosomal protein S6 | RPS6 |
| CENPA_1_14 | MGPRRRSRKPEAPR | Histone H3-like centromeric protein A | CENPA |
| PPR1A_28_40 | QIRRRRPTPATLV | Protein phosphatase 1 regulatory subunit 1A | PPP1R1A |
| RYR1_4317_4329 | VRRLRRLTAREAA | Ryanodine receptor 1 | RYR1 |
| CSF1R_701_713 | NIHLEKKYVRRDS | Macrophage colony-stimulating factor 1 receptor precursor | CSF1R |
| STK6_283_295 | SSRRTTLCGTLDY | Serine/threonine-protein kinase 6 | AURKA |
| ADDB_706_718 | KKKFRTPSFLKKS | Beta-adducib | ADD2 |
| KIF2C_105_118_S106G | EGLRSRSTRMSTVS | Kinesin-like protein KIF2C | KIF2C |
| ERBB2_679_691 | QQKIRKYTMRRLL | Receptor tyrosine-protein kinase erbB-2 precursor | ERBB2 |
| H32_3_18 | RTKQTARKSTGGKAPR | Histone H3.2 | H3C15 |
| TOP2A_1463_1475 | RRKRKPSTSDDSD | DNA topoisomerase 2-alpha | TOP2A |
| GBRB2_427_439 | SRLRRRASQLKIT | Gamma-aminobutyric acid receptor subunit beta-2 precursor | GABRB2 |

D

| Peptide | Sequence | Name | Symbol |
|---|---|---|---|
| ART_025_CX... | GLRRWSLGGLRRWSL | Based on kemptide | PRKX |
| CREB1_126_138 | EILSRRPSYRKIL | cAMP response element-binding protein | CREB1 |
| GBRB2_427_439 | SRLRRRASQLKIT | Gamma-aminobutyric acid receptor subunit beta-2 precursor | GABRB2 |
| MYPC3_268_280 | LSAFRRTSLAGGG | Myosin-binding protein C, cardiac-type | MYBPC3 |
| NCF1_321_333 | QDAYRRNSVRFLQ | Neutrophil cytosol factor 1 | NCF1 |
| RS6_228_240 | IAKRRRLSSLRAS | 40S ribosomal protein S6 | RPS6 |
| RYR1_4317_4329 | VRRLRRLTAREAA | Ryanodine receptor 1 | RYR1 |
| VASP_271_283 | LARRRKATQVGEK | Vasodilator-stimulated phosphoprotein | VASP |

**Figure S2. Structural properties of peptides mapped to AKT1 and AKT3.** Sequence logos show frequency (y-axis) of amino acids by position (x-axis) in low, medium, and high affinity peptides for AKT1 (A) and AKT3 (B). Color shows chemistry of amino acids. Tables show metadata of high affinity peptides for AKT1 (C) and AKT3 (D).
